# Monotremes provide novel insights into evolution of the *DMRT* gene family in vertebrates

**DOI:** 10.64898/2026.07.14.738559

**Authors:** Rachel van der Ploeg, Linda Shearwin-Whyatt, Frank Grützner

## Abstract

*Doublesex* and *mab-3* related (*DMRT*) genes encode a family of transcription factors central to sexual development across metazoa. *DMRT* genes are characterised by a highly conserved DNA binding domain (DM) while flanking regions may vary between species. Gene duplication and loss has shaped the diversity of the DMRT genes with several unresolved questions about their evolution. The most well characterised and conserved *DMRT* gene, *DMRT1*, functions as a sexual regulator universally in metazoans. In chicken, *DMRT1* is located on the Z chromosome and acts as a dosage dependent primary sex determination gene. In therian mammals *DMRT1* is autosomal, however, two copies are required for male development. Interestingly in the basal lineage of egg-laying mammals (monotremes), *DMRT1* is localised on the X specific part of one of the X chromosomes. This provided the first evidence of a sex chromosome system with homology to the avian Z chromosome and raises questions about the function and evolution of *DMRT1* in egg-laying mammals. To gain insight into the evolution of mammalian *DMRT* genes we performed sequence and expression analysis of monotreme *DMRT* genes and comparative analysis with other vertebrates. In monotremes, we identified *DMRT* genes 1-7, and show that *DMRT8* is absent, suggesting that *DMRT8* evolved in therian mammals after the divergence of monotremes. Sequence and expression analysis revealed multiple monotreme specific *DMRT1* isoforms with additional protein-coding exons. The independent evolution of monotreme specific changes in *DMRT1* may be the first indication of functional or regulatory differences in monotreme *DMRT1*.

**Article Summary:** Genes in the *Doublesex* and *mab-3* related (*DMRT*) family play important roles in sexual development across animals, but a comprehensive analysis of these transcription factors is lacking in the most basal mammalian lineage of monotremes. This comparative analysis of *DMRT* genes in monotremes and other vertebrates shows the conservation of *DMRT* genes 1– 7 but found no evidence of *DMRT8* in monotremes or marsupial species, suggesting that this gene evolved in eutherians after the divergence of marsupials. The discovery of several monotreme specific isoforms and novel exons of the X linked *DMRT1* reveals unique evolutionary changes in monotreme *DMRT1*.

## Introduction

The evolutionary conservation of the *Doublesex* and *mab-3* related transcription factor (*DMRT*) gene family in metazoans points to a deeply rooted function in the regulation of sexual development and sexually dimorphic functions. *DMRT* genes have also diversified in function and expression across different tissues and developmental stages. These transcription factors are homologous to *Doublesex* (*dsx*) in *Drosophila melanogaster* and *Male abnormal 3* (*Mab-3)* in *Caenorhabditis elegans,* both of which play central roles in sex determination. *DMRT* genes have been found in all Metazoan lineages, including groups such as Porifera (sponges), indicating that the gene family originated in the common ancestor of eumetazoans (Chong et al. 2013; Riesgo et al. 2014; Wexler et al. 2014). Most animals possess multiple *DMRT* genes, ranging from two in *Xenopus laevis*, to eleven in *Caenorhabditis elegans* (Kopp 2012; Watanabe et al. 2017), with eight in therian mammals (Bellefroid et al., 2013; Veith et al., 2006).

A defining feature of the *DMRT* gene family is the conserved DNA binding DM domain composed of six conserved intertwined cysteines and two histidines that bind DNA in a unique manner (Zhu et al. 2000). Structural studies have shown that the recognition helix makes base-specific contacts with the major groove, while the zinc-binding modules bind to the minor groove through contact with the phosphate backbone (Murphy et al. 2015). Outside the DM domain, sequence conservation of *DMRT* genes is limited across taxa (Bellefroid et al., 2013; Volff et al., 2003).

Eight *DMRT* genes (*DMRT1–8*) have been identified across the mammalian lineage. Functional studies in mice have shown that *DMRT1*, *DMRT4* (*DMRTA1*), *DMRT6* (*DMRTB1*), and *DMRT7* (*DMRTC2*) play key roles in gonadal development, whereas other family members (*DMRT2*, *DMRT3*, and *DMRT5* (*DMRTA2*)) have broader developmental roles (Raymond et al. 2000; Balciuniene et al. 2006; Kawamata and Nishimori 2006; Zhang et al. 2014; Zarkower and Murphy 2022). *DMRT8* lacks the conserved DM domain, and its function remains unknown (Veith et al. 2006). Notably, therian mammals can possess up to four copies of *DMRT8*, all located on the X chromosome (Veith et al. 2006). Despite the absence of a DM domain, *DMRT8* has been hypothesised to have a domain-independent role, potentially acting as a regulator of other *DMRT* proteins during male sexual development (Veith et al. 2006).

*DMRT1* is the most extensively studied *DMRT* gene and exhibits a deeply conserved role in sexual development, specifically in male development and sex determination. In birds, *DMRT1* is located on the Z chromosome and functions as the master sex determining gene (Nanda et al. 1999; Smith et al. 2009; Lambeth et al. 2014). Gene dosage drives testis development in chicken, where expression of two copies of *DMRT1* in ZZ genotype (males) is required to initiate gonad differentiation into testes, via transcriptional activation of *SOX9* and suppression of aromatase expression (Smith et al. 2009). Knockdown of *DMRT1* expression in genetically male chicken embryos results in partial sex reversal, causing feminisation of the gonads (Ayers et al. 2015).

In contrast to birds, *DMRT1* is autosomal in most other metazoans, in humans it is located on the distal short arm of chromosome 9 which has extensive homology to the chicken Z chromosome (Raymond et al. 1998; Nanda et al. 1999). Deletion of the *DMRT1* containing short distal arm of chromosome 9 results in Human XY sex reversal and gonadal dysgenesis (McDonald et al., 1997; Veitia et al., 1997). In humans, *DMRT1* is expressed in male and females during foetal development and two copies of *DMRT1* are required for normal male gonadogenesis (Zarkower and Murphy 2022). *DMRT1* expression in adults is restricted to the testis (Raymond et al. 1998) (Jørgensen et al. 2012). Studies in the mouse show that *DMRT1* plays a role in male, but not female, gonadal development (Zarkower and Murphy 2022). *DMRT1* is required for male somatic and germ cell differentiation and the maintenance of Sertoli cells in the mouse testis (Raymond et al. 1999; Lei et al. 2007; Matson et al. 2010). Mice with a deletion of both *DMRT1* alleles have impaired postnatal testis development (Raymond et al. 2000). Loss of *DMRT1* expression in postnatal testis results in reprogramming of Sertoli cells to granulosa cells (Minkina et al. 2014).

Outside of the amniotes, *DMRT1* plays a central and conserved role in vertebrate sex determination and has repeatedly been co-opted as a master sex determining gene in multiple lineages. The acquisition of novel sex determination mechanisms has been associated with modification or duplication of *DMRT1* in several vertebrate clades (Zarkower and Murphy 2022). Independent duplication and neofunctionalisation events in the genera *Oryzias* and *Xenopus* gave rise to two master sex determining genes, *dmy* and *DM-W*, respectively (Mawaribuchi et al. 2012; Yoshimoto and Ito 2011). In medaka (*Oryzias latipes*), which possesses an XY sex chromosome system, the Y localised gene *DMRT1bY/dmy* originated from duplication of autosomal *DMRT1* and was established as the master sex-determining gene through functional analyses (Matsuda et al. 2002; Matsuda 2005; Nanda et al. 2002). The function of *dmy* is analogous to the mammalian sex determination gene, *SRY* (Nagahama et al. 2021). In contrast, in the African clawed frog (*Xenopus laevis*), which has a ZW sex chromosome system, the W-linked gene *DM-W* arose via partial duplication and C-terminal truncation of *DMRT1*. *DM-W* dominantly inhibits autosomal *DMRT1* function, and this truncated DM protein promotes female development (ovary formation) while supressing testis differentiation (Yoshimoto et al. 2008; Bewick et al. 2011). Recent work identified an amino acid substitution, from serine (S) to threonine (T) at position 15 (S15T) in the DM domain of both *dmy* and *DM-W*, which is under positive selection and enhances DNA binding and transcriptional regulation (Ogita et al. 2020). This convergence supports a common evolutionary mechanism by which *DMRT1* paralogues acquire master sex determining function.

Despite research in therian mammals and non-mammalian vertebrates, comparatively little is known about the evolution of *DMRT* genes in monotremes. The monotreme lineage, which consists of the platypus (*Ornithorhynchus anatinus*) and the echidna (*Tachyglossus aculeatus*), occupy a pivotal phylogenetic position as sister to therian mammals and possess a unique multiple XY sex chromosome system with homology to avian sex chromosomes. Monotremes diverged from therian mammals around 187 million years ago, and the platypus and echidna diverged around 55 million years ago (Zhou et al. 2021). Previous research focusing on monotreme *DMRT1*, mapped this gene to a monotreme X chromosome rather than an autosome (Grützner et al. 2004). This unexpected chromosomal context raises questions about the evolutionary and function in particular of *DMRT1* in monotremes.

Here we present analysis of monotreme *DMRT* genes, which provide novel insights into the lineage specific changes to the *DMRT* gene family. Including the differential retention and loss of *DMRT* genes across vertebrates. The monotreme lineage was found to have seven *DMRT* genes (*DMRT1-7*), and *DMRT8* is not present. Additionally monotreme specific exons and tissue specific splice variants of monotreme *DMRT1* were discovered. Together, these findings provide insights into the evolutionary diversification of the *DMRT* gene family and its potential role in the monotreme lineage.

## Materials and Methods

### Sequence and phylogenetic analysis

DM domain containing genes were identified in the two available Monotreme genomes, *Ornithorhynchus anatinus* (mOrnAna1.pri.v4) and *Tachyglossus aculeatus* (mTacAcu1.pri). A reciprocal BLAST approach was used using the 8 characterised full-length mammalian and chicken *DMRT* proteins, and the DM domain alone were conducted. Monotreme genomes were also searched for genes annotated as DM domain containing. Phylogenetic analyses involved generating an alignment of DM Domain containing amino acid sequences in Geneious Prime 2025.1.2 (https://www.geneious.com) using MUSCLE 5.1 (algorithm PPP) with default settings (Edgar 2022). Accession numbers for amino acid sequences used are documented in Supplementary Table 1. A phylogenetic tree was then constructed using Geneious Prime 2025.1.2 (https://www.geneious.com) from the protein alignment. Jukes-Cantor genetic distance model and UPGMA tree building method were used, (default settings). Tree was resampled via Bootstrap, with 10,000 replicates and a support threshold of 50%.

### Synteny analysis

NCBI Genome data viewer (Rangwala et al. 2021) and Genomicus 2021-08-15 (Nguyen et al. 2022) were used for synteny analysis of monotreme *DMRT1-7* and eutherian *DMRT8*. The position and orientation of these genes were compared within the tetrapods, the chromosomal locations of each also noted (Table S2).

### Transcriptome analysis

Gene and isoform expression levels were quantified from RNA-Seq data using RSEM 1.3.3 (Li and Dewey 2011). RNA-Seq fastq files for platypus and echidna tissues (accessions GSE30352, GSE97367, PRJNA1284983 and PRJNA591380) were mapped to the platypus (mOrnAna1.pri.v4) and short-beaked echidna (mTacAcu1.pri) genomes and normalised TPM and FPKM values were generated. The Human Protein Atlas (proteinatlas.org) and Galbase (Fu et al. 2022) was used to compare transcriptome data from adult human and chicken tissue to the generated monotreme data. Gene expression was visualised using GraphPad Prism version 10.5.0 (673). Integrative Genomics Viewer (IGV) 2.16.0 was used to visualise RNA-Seq reads mapping to the monotreme specific exons in echidna and platypus.

### Reverse transcription polymerase chain reaction

RNA extractions from snap frozen frozen platypus and echidna tissue stored at −80°C were conducted using TRIzol® (Invitrogen) for platypus and NucleoZOL (Machery-Nagel™) for echidna (manufacturer’s instructions followed). Total RNA samples were stored at −80°C after re-suspension in nuclease free water. g-DNA was removed using the DNA-free™ DNA Removal Kit (Invitrogen™) (manufacturer’s instructions followed). Total RNA samples from echidna and platypus were converted to cDNA using iScript™ cDNA Synthesis Kit (Bio-Rad). The cDNA synthesised was stored at −20°C.

Diagnostic primers were designed to amplify regions of the monotreme *DMRT1* gene (primer sequences are included in Table S3). OneTaq® DNA Polymerase (New England Biolabs, Ipswich, MA, USA) was used for polymerase chain reactions. 25µl reactions were prepared containing 2.5µl of 10mM dNTPs, 5µl of OneTaq® Standard Reaction Buffer, 0.2µl of OneTaq® DNA Polymerase, 13.3µl of nuclease-free H_2_O, 1µl each 10µM forward and reverse primers, 2µl of cDNA. PCR cycling conditions initial denaturation at 94.0°C for 3 minutes, followed by 36 cycles of denaturation at 94.0°C for 30 seconds, annealing at 57.0°C for 30 seconds and elongation at 68.0°C for 1 minute, with a final extension at 68.0°C for 5 minutes. PCR products were visualised on a 2% agarose gel stained with SYBR Safe DNA Gel Stain (Invitrogen™).

### Isoform identification through cloning and sequencing of RT-PCR products

PCR products were excised and the DNA extracted using Monarch® DNA Gel Extraction Kit according to manufactures instructions. Purified PCR products were then cloned using the pGEM®-T Easy Vector System (Promega) into NEB® 5-alpha Competent E. coli (High Efficiency). PCR colony screening of products on X-Gal + IPTG treated ampicillin agar plates was completed using OneTaq® DNA Polymerase (New England Biolabs, Ipswich, MA, USA). 25µl reactions were prepared containing 2.5µl of 10mM dNTPs, 5µl of OneTaq® Standard Reaction Buffer, 0.2µl of OneTaq® DNA Polymerase, 10.3µl of nuclease-free H_2_O, 1µl each 10µM forward and reverse primers, 5µl of bacteria colony suspended in nuclease-free H_2_O. PCR cycling conditions initial denaturation at 94.0°C for 3 minutes, followed by 36 cycles of denaturation at 94.0°C for 30 seconds, annealing at 57.0°C for 30 seconds and elongation at 68.0°C for 1 minute, with a final extension at 68.0°C for 5 minutes. PCR products were visualised on a 2% agarose gel stained with SYBR Safe DNA Gel Stain (Invitrogen™). PCR products were then excised and the DNA extracted cleaned up using QIAquick PCR Purification Kit. Sanger sequencing of products was then completed to confirm sequences and *DMRT1* exons identified.

## Results

### Evolution and lineage specific diversification of *DMRT* genes in monotremes and other vertebrates

To examine the evolution of the *DMRT* gene family in mammals, the genes were analysed using the recently published platypus and echidna genomes (Zhou et al. 2021; Zhou et al. 2025). Reciprocal BLAST and synteny analyses identified seven *DMRT* genes in both monotremes (*DMRT1*, *DMRT2*, *DMRT3*, *DMRT4* (*DMRTA1*), *DMRT5* (*DMRTA2*), *DMRT6* (*DMRTB1*), and *DMRT7* (*DMRTC2*)) (Figure 1a, Table S2). In contrast, *DMRT8* (*DMRTC1*), which is present in other mammalian lineages, was absent in monotreme genomes (Figure 1a, Table S2).

**Fig. 1.**
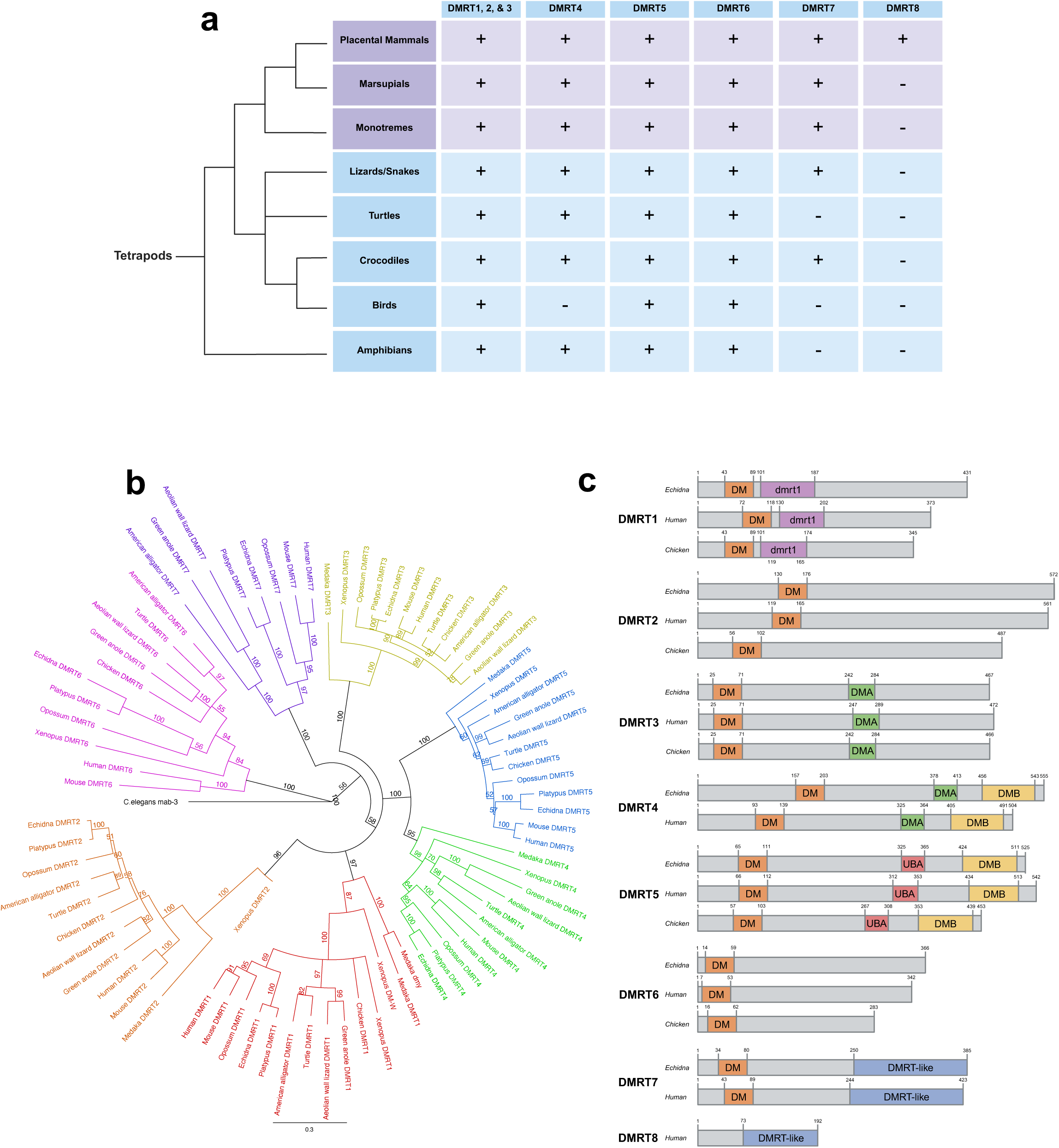
Analysis of *DMRT* gene family members. (a) Summary of *DMRT* gene family members present in various tetrapod clades. (b) Phylogenetic tree from protein alignment of *DMRT* genes. (c) A schematic diagram of conserved domain structures in *DMRT* proteins from Echidna, Human, and Chicken.

Phylogenetic analyses were performed using full length *DMRT* protein sequences from both platypus and echidna, together with representative vertebrate species (Figure 1b). The resulting tree topology was consistent with established species phylogeny, and each *DMRT* gene formed a monophyletic clade, except for *DMRT8*, which clustered within the *DMRT7* lineage (*DMRT* gene phylogeny excluding *DMRT8* shown in Figure 1b; phylogeny including *DMRT8* shown in Figure S1). Within each *DMRT* lineage, platypus and echidna *DMRT* genes formed monophyletic clades with high bootstrap support (Figure 1b) confirming the correct identification of the monotreme *DMRT* genes.

Analysis of amniote *DMRT* proteins revealed conservation of organisation across amniotes and a comparison between echidna, human and chicken are shown in Figure 1c. To investigate the evolution of *DMRT* genes within the monotreme lineage, full length *DMRT* amino acid sequences from the platypus and echidna were compared. In particular, the DNA-binding DM domain was highly conserved between orthologous platypus and echidna *DMRT* proteins, with greater than 90% sequence identity (Table S4). In contrast, conservation across the full length *DMRT* proteins was lower, ranging from 55–97% sequence identity, as has previously been noted for *DMRT* genes (Figure S2, Table S4).

Comparative synteny analysis was carried out to confirm identity and genomic organisation of monotreme *DMRT* genes. In all species *DMRT1-3* genes are located together in a highly conserved genomic region. Previously it was shown that *DMRT1-3* are located adjacently on the platypus X5 sex chromosome (El-Mogharbel et al. 2007). With the echidna genome available and an improved platypus genome we determined that *DMRT1-3* occupies the same genomic region in platypus and echidna, with echidna *DMRT1-3* located on the X4 sex chromosome, which shares homology to platypus X5. Synteny analysis confirmed that the order and orientation of *DMRT1–3*, as well as the flanking genes *DOCK8*, *KANK1*, *SMARCA2*, and *VLDLR*, are conserved across all tetrapod species analysed (Figure 2a). However, lineage specific gene insertions in both platypus and echidna disrupt collinearity within this region (Figure 2a).

**Fig. 2.**
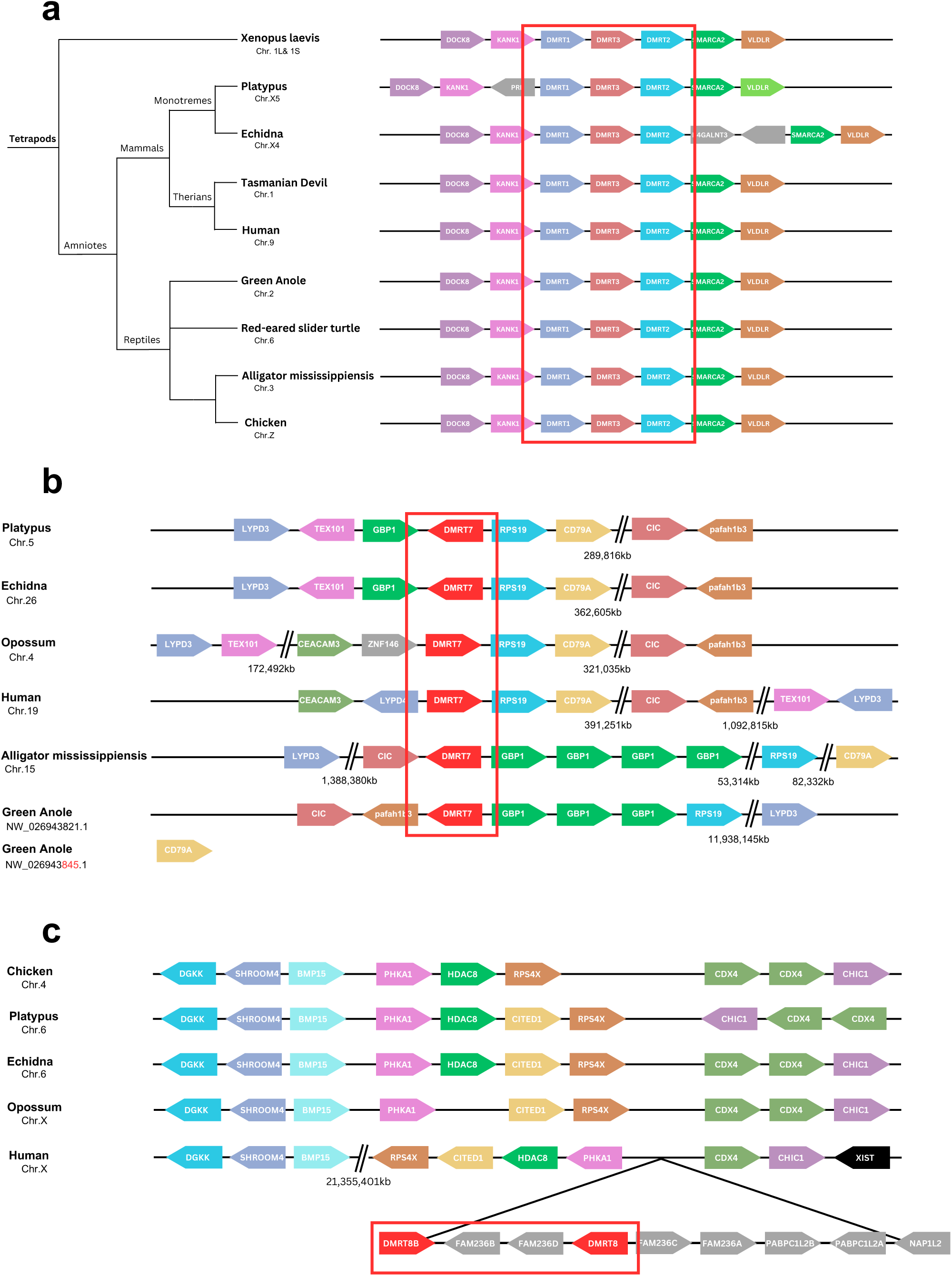
Synteny analysis of *DMRT* gene family members. (a) *DMRT1*, *DMRT2*, and *DMRT3*, unnamed grey box is uncharacterized LOC119947860 (b) *DMRT7*. (c) *DMRT8*.

*DMRT4* is not linked to the *DMRT1-2-3* cluster, it is located further upstream on the same chromosome in both monotreme species (echidna X4, and platypus X5), as well as in all other tetrapod species analysed, with the exception of avian genomes (*G. gallus*, *T. guttata*, & *P. domesticus*), in which *DMRT4* orthologues were not identified (Figure 1a, Figure S3, Table S2). Synteny analysis indicated that the genomic region containing *DMRT4* in other tetrapods is missing in the chicken as representative of avian genomes (Figure S3).

In both echidna and platypus, *DMRT5* was located on chromosome 18, with conserved synteny across tetrapods (Figure S4). In contrast, *DMRT6* was located on different chromosomes in echidna (chromosome 10) and platypus (chromosome 18), and synteny was not conserved in monotremes compared to other species analysed (Figure S5). In monotremes, the orientation of *DMRT6* relative to the flanking genes has changed, suggestive of an inversion of the gene (Figure S5). In addition, a synteny break has separated genes (*LRP8* and *MAGOH*) downstream of *DMRT6* onto different chromosomes in platypus and echidna (chromosome 17 and 24, respectively) (Figure S5). Instead, a single annotated gene, *TEX47*, is located downstream of *DMRT6* in monotremes, at the end of the chromosome. *TEX47* is not syntenic with *DMRT6* in other species, for example, in humans, *DMRT6* is located on chromosome 9, whereas *TEX47* is in a different genomic region on chromosome 7 (Figure S5).

*DMRT7* has previously been reported as a mammalian specific gene involved in spermatogenesis (Kim et al. 2007; Tsend-Ayush et al. 2009a). *DMRT7* was identified on platypus chromosome 5 and echidna chromosome 26 and we also identified orthologues in recently published reptilian genomes for the first time (*A.* carolinensis, *A. mississippiensis*, *P. raffonei*, & *C. aspera*) (Figure 1a, Figure 2b). *DMRT7* synteny is conserved among mammals and partially conserved in the reptiles (Figure 2b). Only a subset (*GBP1* and *RPS19*) of mammalian *DMRT7* flanking genes are conserved in the immediate flanking regions of *A.mississippienis* (American alligator) and *A.carolinensis* (Green anole) (Figure 2a).

*DMRT8* has also been reported as mammalian specific, however, this gene was absent from both monotreme genomes and from the investigated marsupial species (*Trichosurus vulpecula*, *Sarcophilus harrisii*, *Monodelphis domestica*, & *Antechinus flavipes*), suggesting that it is in fact eutherian mammal specific (Figure 1a and Table S2). *DMRT8* copy number varied among eutherian mammals, with one copy in *Macaca mulatta*, two in *Homo sapiens*, three in *Mus musculus*, and four in *Rattus norvegicus*. Synteny analysis further supports the absence of *DMRT8* outside of eutherian mammals. The genomic region upstream of *DMRT8* is conserved across amniotes (Figure 2c), whereas downstream flanking genes in humans (*RPS4X, CITED1, HDAC8* and *PHKA1*), retain synteny but not collinearity across amniotes (Figure 2c). Additional downstream synteny breaks were observed between humans and other amniote species. This genomic region is autosomal in all species analysed, except in eutherian mammals where it is located on the X sex chromosome.

### Comparative analysis of *DMRT* gene expression in monotremes and other amniotes

*DMRT* gene expression was compared across a range of adult tissues in humans, monotremes, and chicken using available transcriptome data. Platypus and echidna exhibited broadly similar *DMRT* gene expression profiles, with most *DMRT* genes expressed in the adult testis (Figure 3a & 3b, Table S5). All seven *DMRT* genes were expressed in platypus testis, whereas all except *DMRT5* were expressed in echidna testis. In humans, all *DMRT* genes were expressed in adult testis except *DMRT4*, which is consistent with previous studies, where RT-PCR showed expression of all *DMRT* genes (*DMRT1-8*) in mature mouse testis (Figure 3c, Table S5) (Balciuniene et al., 2006; Kim et al., 2003; Ono et al., 2021; Veith et al., 2006). In chicken testis, four of the five chicken *DMRT* genes were expressed, *DMRT3* was not expressed, and it is also not expressed during chicken testis development (Smith et al. 2002)) (Figure 3d, Table S5).

**Fig. 3.**
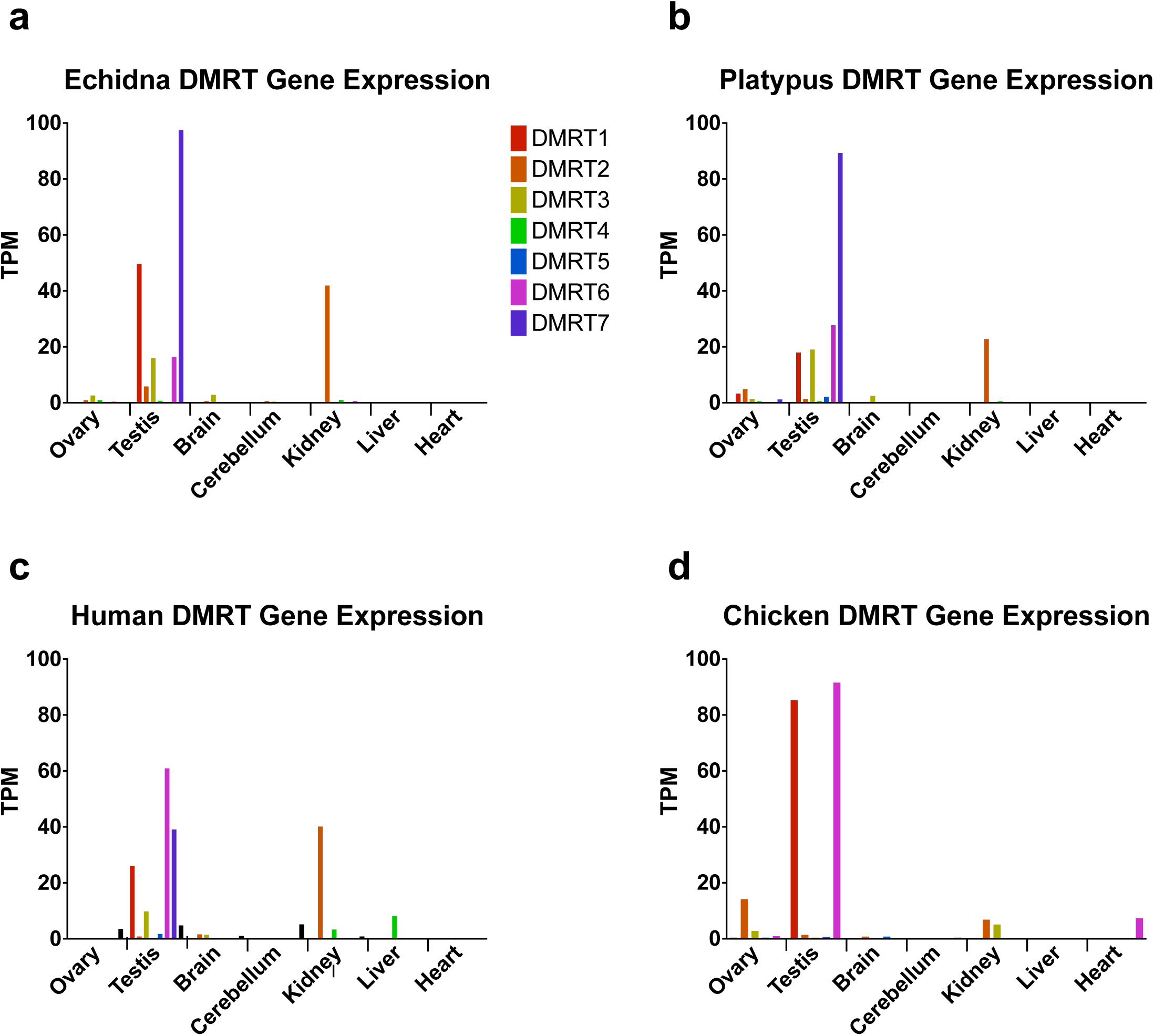
DMRT gene family expression from RNA-seq data. *DMRT* gene family expression in (a) Echidna, (b) Platypus, (c) Human, (d) Chicken. RNA-Seq fastq files for platypus and echidna tissues were mapped to platypus (mOrnAna1.pri.v4) and short-beaked echidna (mTacAcu1.pri) genomes and RSEM 1.3.3 was used to estimate gene expression levels. Monotreme nTPM values were then visualised and The Human Protein Atlas (proteinatlas.org) and Galbase (Fu et al. 2022) were used for human and chicken nTPM values.

In contrast, *DMRT* gene expression in adult ovary differed among species. In humans, only *DMRT8* was expressed in adult ovary. This contrasts with monotremes and chicken, in which multiple *DMRT* genes were expressed, albeit at low levels. In platypus ovary, expression of *DMRT1*, *DMRT2*, *DMRT3*, *DMRT4*, and *DMRT7* was detected, whereas echidna ovary expressed *DMRT2*, *DMRT3*, and *DMRT4* (Figure 3a & b, Table S5). A similar pattern was observed in chicken ovary, where *DMRT2*, *DMRT3*, and *DMRT6* were expressed, highlighting a broader ovarian *DMRT* expression profile outside eutherian mammals.

*DMRT1* is the most extensively studied member of the *DMRT* gene family and has been shown to be expressed in testis across numerous species during development and adulthood (Kim et al. 2003; Tsend-Ayush et al. 2009b). Consistent with these findings, *DMRT1* expression was detected in adult platypus and echidna testis. Notably, *DMRT1* expression was also observed in the adult platypus ovary but was absent from the echidna ovary (Figure 3b, Table S5).

### Discovery of novel exons in the monotreme *DMRT1* gene

Alternative splicing of *DMRT1* transcripts has been observed during development, including during sex determination in testis (Augstenová and Ma 2025). Across vertebrates, *DMRT1* has five highly conserved protein coding exons, based on predicted *DMRT1* genes across taxa (Figure 4a). Outside of the DM domain, *DMRT1* protein sequences were not well conserved across vertebrates (Figure S6) Genome annotations suggested the presence of multiple *DMRT1* isoforms in monotremes (Figure 4a). The number of predicted *DMRT1* isoforms differed between monotreme species, with four predicted isoforms in echidna and one in platypus.

**Fig. 4.**
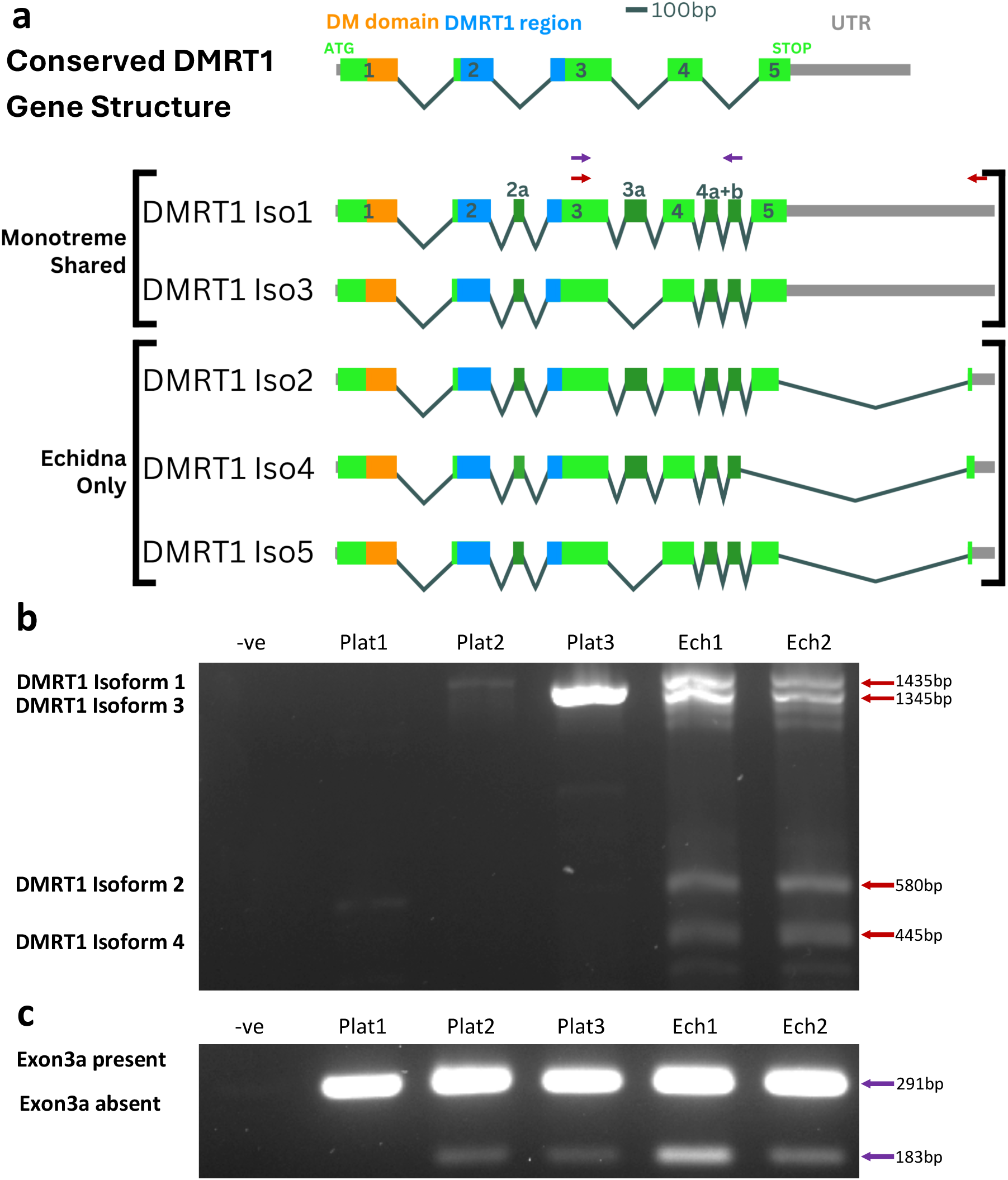
DMRT1 isoform analysis. (a) Conserved *DMRT1* transcript structure compared with monotreme *DMRT1* transcript structures is shown. Exons are drawn to scale, where 1-5 are the conserved exons and 2a, 3a, 4a+b are the novel monotreme specific exons. The grey bar shows the untranslated region (UTR), the two conserved regions, DM domain and *DMRT1* region are highlighted. RT-PCR to confirm expression of different monotreme *DMRT1* transcripts was performed with diagnostic primers (red arrows indicate primers for RT-PCR in (b), purple arrows indicate primers for RT-PCR in (c)), using cDNA from both platypus and echidna adult male testis tissue (b) and (c). (b) expression of different monotreme *DMRT1* isoforms (red arrows) expected product sizes: isoform 1: 1435bp, isoform 3: i345bp, isoform 2: 580bp, isoform 4: 445bp. (c) expression of *DMRT1* transcripts indicating the presence (291bp) or absence (183bp) of exon3a (purple arrows).

Analysis of the predicted monotreme *DMRT1* isoforms suggested that they carried additional coding exons (called 2a, 3a, 4a and 4b, Figure 4a). To assess expression of these isoforms, RT-PCR and sequencing was performed using testis RNA. This confirmed expression of multiple *DMRT1* isoforms in monotreme testis, with four transcripts detected and confirmed with sequencing in echidna and two in platypus (Figure 4b). Due to the limited availability of platypus tissue, we were unable to confirm with sequencing if all echidna transcripts were also expressed in platypus testis. Analysis of monotreme *DMRT1* transcripts revealed that the four novel exons (2a, 3a, 4a, and 4b), are non-homologous to the five conserved exons (1–5) (Figure 4a). Exons 2a, 4a, and 4b were predicted in both monotreme species. Exon 3a was predicted only in the echidna. Expression of exon 2a, 4a and 4b, as well as exon 3a in both species was confirmed by RT-PCR and sequencing (Figure 4, Figure S7). The sequences of these novel exons are not homologous to the ancestral protein coding regions (Table S6 & Table S7). Transcriptome data was used to count RNA-Seq reads mapping to the monotreme specific exons in echidna and platypus (Table S8).

To compare the evolution of these novel protein coding regions within the monotreme lineage, protein sequences of the monotreme novel exons were compared between echidna and platypus. Sequencing confirmed exon 3a was mutation free, whereas 2a had one conservative amino acid change between platypus and echidna (S to A) (Figure S7a+b). Exon 4a and 4b were unable to be sequenced in the platypus. The NCBI predicted sequence of exon 4a and 4b in the platypus was compared to the confirmed echidna sequences. Exon 4a and 4b each had three amino acid changes between platypus and echidna (Figure S7c). All three amino acid substitutions on exon 4a were non-conservative, and all three substitutions on exon 4b were conservative (Figure S7c).

Phylogenetic analysis of both *DMRT1* full length proteins (Figure S8a and S8c) and *DMRT1* proteins excluding the highly conserved DM domain (excluding exon 1) (Figure S8b), was conducted to analyse the evolution of the monotreme novel exons. Platypus and echidna *DMRT1* sequences branched outside of all other vertebrate species analysed (Figure S8).

The monotreme specific protein coding exon2a is located within the conserved *dmrt1* domain (Figure 4a). In monotremes, the amino acid residues flanking exon 2a are of particular interest due to their previously reported association with modes of sex determination in reptiles (Figure S9) (Janes et al. 2014). Specifically, changes to the amino acids at positions T54 and S57 have been correlated with reptile species having either temperature-dependent or genetic sex determination (Janes et al. 2014). This association has not been observed in mammals as there is no temperature-dependent sex determination (Janes et al. 2014). A difference was observed between monotreme species at these positions, where echidna retained the ancestral threonine (T) and serine (S) residues, whereas in platypus the serine (S) residue was replaced by alanine (A) (Figure S9).

## Discussion

The *DMRT* gene family is deeply conserved across metazoans and plays fundamental roles in development, most prominently in sex determination and sexual differentiation. Here, we provide a comprehensive analysis of the identity, organisation, and evolution of *DMRT* genes in monotremes and compare them with key vertebrate lineages.

### *DMRT* gene family conservation and gene loss

Genome analyses revealed strong conservation of *DMRT1-6* genes across tetrapods, except for *DMRT4* in birds. *DMRT4* was not identified in avian genomes (*Gallus gallus*, *Taeniopygia guttata*, and *Passer domesticus*) and was likely lost after divergence from crocodilians approximately 230-240 million years ago (Green et al. 2014; Brusatte et al. 2015). In other vertebrates, *DMRT4* has roles in sex differentiation, neurogenesis, and olfactory system development (Bellefroid et al., 2013; Hong et al., 2007; Huang et al., 2005), and in mice *DMRT4* contributes to folliculogenesis and male sexual behaviour (Balciuniene et al., 2006). It is unclear if other *DMRT* genes have adopted these functions in avian lineages in the absence of *DMRT4*.

Monotremes shared all *DMRT* genes (*DMRT1-7*) with eutherian mammals, except *DMRT8* (*DMRTC1*) (Figure 1a, Table S2). *DMRT8* was also missing in marsupial genomes (Figure 1a, Figure 2c, and Table S2). This challenges earlier reports proposing an early mammalian origin (Veith et al. 2006) and instead supports a more recent emergence in eutherians (Luo et al. 2011), likely via a duplication of *DMRT7* onto the eutherian X chromosome (Ottolenghi et al. 2002; Veith et al. 2006).

### Redefining the origin of *DMRT7* and *DMRT8*

*DMRT7* has previously been reported as a mammalian specific gene (Kawamata and Nishimori 2006; Veith et al. 2006; Kim et al. 2007; Kawamata et al. 2008; Tsend-Ayush et al. 2009b; Zhang and Zarkower 2017). *DMRT7* orthologues were identified in monotremes confirming previous reports. However, we identified *DMRT7* in the genomes of both the American alligator (*Alligator mississippiensis*) and the green anole (*Anolis carolinensis*) (Figure 1a, Table S2 and Figure 2b). This suggests that *DMRT7* is not mammalian specific and evolved earlier in vertebrates.

In therian mammals, *DMRT7* is essential for spermatogenesis, localises to the XY body during meiosis, and is required for the transition between meiotic sex chromosome inactivation (MSCI) and post-meiotic sex chromatin (PMSC). The evolution of *DMRT7* in the mammal lineage alongside therian sex chromosomes was therefore thought to be linked with MSCI evolution (Kawamata and Nishimori 2006; Veith et al. 2006; Kim et al. 2007; Kawamata et al. 2008; Tsend-Ayush et al. 2009b; Zhang and Zarkower 2017). The presence of *DMRT7* in reptiles challenges this idea. In therian mammals, MSCI occurs during meiotic prophase I and results in transcriptional silencing of heteromorphic sex chromosomes (Turner 2007; Daish et al. 2015). MSCI is common to mammalian sex chromosome systems, occurring in both eutherian mammals and in monotremes, thus it has thought to be an ancestral mammalian process (Murat et al. 2023). In bird lineages there is conflicting evidence, *DMRT7* is missing in the chicken genome, and research suggests MSCI may be lacking or is transient (Schoenmakers et al. 2009a; Guioli et al. 2012). MSCI is also absent in teleost fish, which suggests that sex chromosome specific silencing originated evolved in mammals (Schoenmakers et al. 2009b; Guioli et al. 2012; Daish et al. 2015; Shaw et al. 2024 Nov 29). As *DMRT7* helps regulate the transition from MSCI to PMSC, finding *DMRT7* outside of the mammalian lineage suggests that *DMRT7* originally had a different function and evolved a role in the mammalian lineage. Future work is required to establish a function of *DMRT7* in reptiles.

The origin of *DMRT8* also needs to be redefined as this gene was absent from both monotreme genomes and from the marsupial species *Trichosurus vulpecula*, *Sarcophilus harrisii*, *Monodelphis domestica*, & *Antechinus flavipes* (Figure 1a, Table S2). The absence from mammal lineages outside of eutherians indicates that *DMRT8* is not a universal mammalian gene, but instead likely originated within the eutherian lineage. Phylogenetic analyses placed eutherian *DMRT8* within the *DMRT7* clade, supporting its emergence through duplication of *DMRT7* after divergence of eutherians from other mammalian lineages 160 million years ago (Luo et al. 2011). Broader taxonomic sampling to include other non-eutherian mammals will refine this view.

### Genomic context and *DMRT* gene evolution

Chromosomal localisation of *DMRT* genes differs across vertebrate lineages, and these genes can be found on both autosomes and sex chromosomes. This lineage specific localisation raises questions about how genomic context may influence sequence divergence. Genes located on sex chromosomes are generally reported to evolve more rapidly than autosomal genes (Mank et al. 2010; Kousathanas et al. 2014; Charlesworth et al. 2018). There is also evidence that expression levels of X-linked genes often show faster evolutionary change compared to autosomal genes, potentially reflecting sex specific selective pressures (Brawand et al. 2011; Kayserili et al. 2012; Meisel et al. 2012).

*DMRT1* is autosomal in therian mammals and most other amniotes but is located on sex chromosomes in several other vertebrate lineages, including birds and monotremes. In addition, *DMRT8* is X-linked in eutherian mammals. To test whether chromosomal context influences evolutionary rate (branch lengths), the phylogeny of the *DMRT* gene family across vertebrates was examined. Phylogenetic reconstruction allowed identification of potential lineage specific shifts in gene function or expression. It also provided validation of *DMRT* gene identification in monotremes, as each monotreme *DMRT* protein clustered into the corresponding clades (Figure 1b). In mammals, *DMRT* genes located on sex chromosomes showed increased sequence divergence when compared to *DMRT* genes located on autosomes (Figure S1, and S8). Phylogenetic analysis showed *DMRT8* had significantly longer branch lengths when compared to other *DMRT* genes within the vertebrates (Figure S1). Phylogeny of *DMRT1* proteins showed that the platypus and echidna *DMRT1* sequences have undergone significant evolution (Figure S8). The platypus and echidna *DMRT1* protein branches sit outside of the *DMRT1* vertebrate clade. This was the case for both DMRT1 full length proteins (Figure S8a and S8c) and DMRT1 proteins excluding the highly conserved DM domain (excluding exon 1) (Figure S8b). The variation in chromosomal location across lineages, showed clear association between sex linkage and increased sequence divergence within the *DMRT* family. Genes located on sex chromosomes exhibited consistently elevated evolutionary rates relative to autosomal counterparts (Figure S1, and S8). When platypus and echidna *DMRT* proteins were compared with each other, they formed strongly supported monophyletic relationships, indicating that neither monotreme lineage has experienced substantial divergence of *DMRT* genes since their split.

Comparative analysis of the *DMRT1* DNA-binding domain (DM domain) revealed strong amino acid conservation between the monotremes, and with other vertebrate lineages (Table S4, Figure S6). The high conservation of the *DMRT1* DM domain, suggests that the monotreme *DMRT1* protein binds to target sequences in a similar way to other species (Murphy et al. 2015). This also suggests that ancestral *DMRT1* target genes and function are possibly conserved in the monotreme lineage. Identifying the timing and tissue specificity of *DMRT1* expression in monotremes provides a better indication of potential functional changes.

### Monotreme *DMRT1* is expressed in adult ovary

To examine gene expression changes in the *DMRT* family, newly available monotreme transcriptome information was compared to human and chicken transcriptome data. Monotremes share a similar expression pattern of *DMRT* genes, compared to chicken and human data, suggesting a conserved function of these genes in monotremes (Figure 3, Table S5). *DMRT* genes were shown to have a male biased tissue expression pattern, with expression primarily occurring in testis (Figure 3, Table S5). *DMRT1* was expressed in both adult platypus and echidna testis tissue which supports previous findings. However, RNA-seq data showed *DMRT1* expression in the adult platypus ovary, a pattern not reported in other amniotes, where *DMRT1* is typically restricted to male gonadal tissue. Conservation of gene expression within the same cell type across species generally reflects conserved function. Ovarian *DMRT1* expression has not been reported in vertebrates until, recent work in *Xenopus*, where *DMRT1* was shown to be necessary for female fertility and egg production (Kukoly et al. 2026). A genome duplication of *Xenopus laevis* resulted in two copies of *DMRT1*, and *DMRT1* knockouts showed *DMRT1.S* was required for male fertility and spermatogenesis and *DMRT1.L* was required for female fertility and oogenesis (Kukoly et al. 2026). Given the expression of platypus *DMRT1* in both sexes, sex specific functional evolution of monotreme *DMRT1* should be investigated further.

*DMRT1* expression in the mammalian (mouse) ovary, is inhibited by *FOXL2*. *DMRT1* and *FOXL2* have an antagonistic relationship where *DMRT1* expression in the ovary is inhibited by *FOXL2*, and in testis *DMRT1* silences *FOXL2* expression (Lindeman et al. 2015). A key event during mammalian sexual development is differentiation of Sertoli cells in males and granulosa cells in females. Sertoli vs. granulosa cell fate can be reversed by the loss of a transcription factor (*DMRT1* in males, *FOXL2* in females), which can trigger direct transdifferentation between the two cell types (Uhlenhaut et al. 2009; Matson et al. 2011). Previous research has shown that ectopic expression of *DMRT1* in mammalian ovary leads to Sertoli cell like differentiation *in vivo* (Lindeman et al. 2015). The reprogramming of granulosa cells into Sertoli-like cells has not be observed *in vivo* with other testicular transcription factors (*SOX9*, *GATA4*, *NR5A1/SF1*, and *WT1*) (Buganim et al. 2012). The expression of *DMRT1* in the Platypus ovary therefore could suggest a lineage specific shift in function, potentially reflecting either gain or modification of its canonical role in platypus.

### Monotreme specific novel DMRT1 exons

Alternative splicing of *DMRT1* is common across vertebrates (Mizoguchi and Valenzuela 2020; Augstenová and Ma 2025). The *dsx* gene, the invertebrate homologue of *DMRT1*, has been shown to have sex specific alternate splicing of the transcript (Baker and Ridge 1980; Uhlenhaut et al. 2009; Matson et al. 2011). Both sex specific transcription factors, *dsxM* and *dsxF* have homologous DNA binding domains at their N-terminals, however, they have different target genes due to distinctly different the C-terminal domains (Zhu et al. 2000).

There are five conserved *DMRT1* protein coding exons among vertebrates (1, 2, 3, 4, and 5) (Figure 4). Conserved exons 1-5 were identified in both monotreme species; however, we identified four additional novel protein coding exons (2a, 3a, 4a, and 4b) (Figure 4). These additional exons alter the protein sequence and potentially have functional consequences. The monotreme specific exons were found outside the DM domain in protein coding regions. The DM domain of *DMRT1* in eutherian mammals (human and mouse) has the ability to bind to DNA with multiple stoichiometries (Murphy et al. 2015), however how the rest of the protein interacts with regulatory elements is not well understood. There is no structure of the *DMRT1* protein outside of the DM domain, and it is unclear how these additional protein coding exons may affect the protein structure. As regions outside of the DM domain would have many different functions such as the recruitment of co-activators and co-repressors, we can conclude the addition of these monotreme specific exons may affect protein structure and likely would have some functional importance. While alternative exons were predicted in the echidna genome, none were annotated in the platypus genome. RT-PCR and sequencing showed that echidna and platypus express multiple *DMRT1* transcripts in adult testis tissue, confirming the novel protein coding exons were expressed in both monotremes. Sequencing allowed us to identify which protein coding exons were spliced in each transcript and identify exon skipping in platypus and echidna *DMRT1*.

Additional *DMRT1* protein coding exons have not been observed in species outside of the monotremes. There have been instances of duplication events of existing exons in species (Cross et al. 2020), however the sequences of these novel exons in monotremes are not homologous to the ancestral protein coding regions (Table S6 & Table S7). These four additional novel protein coding exons have likely evolved in the ancestor of platypus and echidna around 55 million years ago. This suggests that the *DMRT1* protein has undergone independent evolution in the monotreme lineage.

The sequence evolution of these monotreme specific exons was investigated through phylogenetic analysis of full length *DMRT1* proteins, and the C terminus (excluding the DM domain). This analysis allowed us to specifically look at the C terminus of the *DMRT1* protein which is where the addition of all 5 of the monotreme specific exons has occurred. The result showed that monotreme *DMRT1* protein branches are located outside of all other vertebrates analysed (Figure S8) due to the monotreme specific exons.

Protein sequences of the monotreme novel exons were compared between echidna and platypus. Exon3a is identical between platypus and echidna whereas, exon2a had one conservative amino acid change between platypus and echidna (S to A) (Figure S7a). Exon 4a and 4b were less conserved between platypus and echidna, with three amino acid changes in each exon (Figure S7b). Conservation of these novel exons between platypus and echidna indicates that these exons were acquired before the divergence of platypus and echidna and the sequence conservation for over 55 million years suggests functional importance.

In reptiles there is evidence that temperature dependent sex determination may involve amino acid shifts in *DMRT1*. A region in the *DMRT1* protein has been shown to have a role in switching between genetic and temperature dependent sex determination in reptiles. The combination of an amino acids pair at the 3’ end of exon 2 in *DMRT1* has been shown to correlate with either temperature sex determination (TSD) or genetic sex determination (GSD) (Janes et al. 2014). T54-S57 is the ancestral state, which correlates with temperature sensitive sex determination in reptiles, and any deviations correlate with genetic sex determination (Figure S9) (Janes et al. 2014).

In monotremes T54-S57 are on either side of one of the monotreme specific exon, exon 2a (Figure S9). Exon 2a, is of particular interest as it is located within the *dmrt1* region, a conserved region of the gene (Figure 4a) (Figure S9) (Huang et al., 2025). The protein coding exon insertion between these two residues in monotremes would change the protein structure in this region and potentially function. However, there is no correlation between TSD and GSD with mutations to T54-S57 in therian mammals (Janes et al. 2014).

Monotremes have a genetic sex determination system and evolved their sex chromosomes system independent to other mammals. The monotreme sexual development pathway is poorly characterised, however there is now strong support that the Y-localized anti-Müllerian hormone gene (*AMHY*) is the primary male sex determination gene and *DMRT1* might be involved in its regulation (Shearwin-Whyatt et al. 2025). *DMRT1* was shown to have a male biased expression in the bipotential foetal echidna gonad from the earliest developmental stage examined, alongside *GATA1* and *AMHY* (Shearwin-Whyatt et al. 2025). In addition, conserved potential binding sites for *DMRT1* were found in the promoters of monotreme *AMHX* and *AMHY*, which suggests that *DMRT1* may be involved in the regulation of *AMHY* expression during monotreme gonad differentiation (Shearwin-Whyatt et al. 2025). While *DMRT1* is located on an X chromosome a Y copy DM gene has never been identified and with the strong support that *AMHY* is the sex determination gene, the acquisition of these novel exons and how they alter the function of the *DMRT1* protein should be explored further.

In summary we have conducted a comprehensive analysis on the *DMRT* gene family in monotreme species (*Ornithorhynchus anatinus* and *Tachyglossus aculeatus*) in comparison with other vertebrate species which provided novel insights into the evolution of this gene family in vertebrates. Our results contradict previous reports showing that *DMRT8* evolved in eutherian species and *DMRT7* evolved much earlier in vertebrates than previously proposed. We also discovered that *DMRT4* has been lost in the avian lineage. RNA-seq data showed male expression bias of *DMRT* genes in monotremes and that *DMRT1* is expressed in the monotreme testis and platypus ovary, but not echidna ovary. The highly conserved gene structure including 5 protein coding exons is altered in the monotreme ancestor with new novel exons, which suggests lineage specific evolution of this gene in monotreme species which may indicate new function in monotremes. Detailed analysis of monotreme *DMRT1* showed novel coding exons which are expressed in the testis.

## Data availability statement

The data underlying this article are available in the article and in its online supplementary material.

## Acknowledgements

We would like to thank Zhipeng Qu for his assistance with the Transcriptome data analysis which was previously published (Dann et al. 2024; Wilson et al. 2025).

## Funding

This work was supported by the Australian Research Council Discovery Project grant to FG (DP110105396). RV was supported by a University of Adelaide Research Scholarship.

## Conflict of Interest

The authors declare no conflict of interest.

## Author Contributions

FG, LS-W, RV experimental design. RV performed experiments. FG provided samples. RV wrote the manuscript, edited by FG and LS-W.

